# When do big problems far away become smaller than the problems closer to home

**DOI:** 10.1101/544593

**Authors:** Tjibbe Donker, Katie L. Hopkins, Susan Hopkins, Berit Muller-Pebody, Tim E.A. Peto, Alan P. Johnson, Neil Woodford, Derrick W. Crook, A. Sarah Walker, Julie V. Robotham

**Affiliations:** The National Institute for Health Research (NIHR) Health Protection Research Unit in Healthcare Associated Infections and Antimicrobial Resistance, University of Oxford, Oxford, UK; Nuffield Department of Medicine, University of Oxford, Oxford, UK; National Infection Service, Public Health England, Colindale, London, UK; NIHR Biomedical Research Centre, Oxford, UK

## Abstract

Infection prevention and control strategies aimed at reducing the occurrence of Carbapenemase-Producing Enterobacteriaceae (CPE) and other antimicrobial-resistant organisms often include advice about screening patients coming from hospitals with a known resistance problem, to prevent introductions into new hospitals by shared patients. We argue that, despite being an efficient method of identifying cases, admission screening for introduction prevention is only effective if the absolute number of imported cases from other hospitals outnumbers the cases coming from the hospital’s own patient population, and therefore is only a feasible control strategy during the start of an epidemic.

## The increasing threat of CPE

An increasing number of control strategies against Carbapenemase-Producing Enterobacteriaceae (CPE) and other antimicrobial-resistant organisms include a focus on coordinating control efforts among hospitals nationally [1] or regionally [2,3]. The need for coordinated control stems from the fact that no single hospital is completely segregated from others: they are all connected by the patients they share, thus creating one large national hospital network. For numerous reasons, some patients need to receive care in different hospitals, for instance because the necessary treatment is not available in the original hospital, or because care can be offered more efficiently elsewhere. These shared patients offer the opportunity for CPE and other organisms to be transmitted from one hospital to another [4].

Patient sharing between hospitals has a strong regional tendency [5], driven by movement patterns to and from large tertiary care centres, such as university hospitals. Often, multiple general hospitals refer patients to the same tertiary care centre for advanced care, connecting all hospitals served by this centre into a single group. These groups tend to fall into clear geographical regions, because referral choice is related to the geographical distance between hospitals. Within these regions, hospitals can quickly share each other’s CPE problem(s), with an outbreak in one hospital easily spreading to the others.

By coordinating infection prevention and control efforts among hospitals that are easily affected by each other’s CPE problems, any outbreak can be approached as a problem of all hospitals combined, avoiding any hospitals being caught off-guard by CPE-positive patients. An example of such a coordinated national response, which included central reporting of detected colonised and infected individuals, contained an outbreak of CPE in Israel after local, single hospital-based interventions had failed [6]. Although the described response was expensive and relied on strict compliance with testing and isolation guidelines in all hospitals, it does show the clear benefit of coordinated efforts among hospitals over approaches that are aimed at individual hospitals.

## Preventing introductions between hospitals

The CPE acute Trust toolkit developed by Public Health England in 2013 [7] reiterates the importance of shared patients in the epidemiology of CPE, as it advises (among other things) hospitals to screen patients with a recent history of admission to a hospital with a ‘known CPE problem’. In this way, it aims to prevent CPE introductions from the ‘problem’ hospital to others. Although the toolkit is designed for individual hospitals i.e. it does not directly promote a coordinated approach between hospitals, it does acknowledge the risk of introductions from other hospitals. By correctly defining which hospitals pose the greatest risks, this should theoretically help to prevent spread from one hospital to the others.

However, the strong regional tendency in shared patients means that some hospitals will need to screen many more patients than others, based on the fact that they share many more patients with a ‘problem’ hospital. This can cause considerable strain on a hospital’s resources, as the toolkit advises hospitals to isolate suspected patients until three consecutive rectal swabs (taken 48 hours apart) have tested negative for CPE, meaning that each recently admitted patient previously discharged from the ‘problem’ hospital will need to be isolated for at least four days. This makes strict implementation of the toolkit harder, if not impossible, for some hospitals [8].

The differences in numbers of patients received from other hospitals (i.e. admitted having previously been discharged from elsewhere) also means that the risk posed by an outbreak in one hospital, expressed as the absolute number of expected introductions, differs greatly. A large problem in a hospital far away in the hospital network can therefore pose a smaller absolute risk of introduction than a small problem in a hospital closer to home [9]. Thus, the focus on hospitals far away with known CPE problems may actually undermine control efforts, because the small problems closer to home get overlooked.

Furthermore, strategies aimed at preventing introductions from one population to the other only delay the epidemic spread [10], because it is impossible to identify all arriving cases. Eventually, CPE will pass through the barriers imposed by import screening, and start spreading through the new population, unless control measures are in place to prevent the endogenous spread. After successful introduction, sustained transmission within the hospital’s own patient population may quickly create a much larger problem than introductions from elsewhere.

## The importation tipping point

Within the United Kingdom, and England specifically, there has been a sharp rise in the number of confirmed CPE isolates over the last decade [9]. This increase is in part driven by an outbreak in the North-West of England, the greater-Manchester area, of primarily KPC-positive CPE. Hospitals in this region are therefore flagged as ‘problem’ hospitals, and other hospitals are advised to screen patients who have been admitted previously to hospitals in the Manchester region. However, over the last 5 years, an increasing number of isolates have been submitted to PHE’s national reference laboratory from hospitals outside the Manchester region (Figure 1). These isolates are not carrying exclusively KPC genes, but also harbour different carbapenemases (primarily OXA-48, NDM, and VIM). They are therefore unlikely to have been imported from hospitals in the Manchester region and signify unrelated spread and transmission.

**Figure 1).**
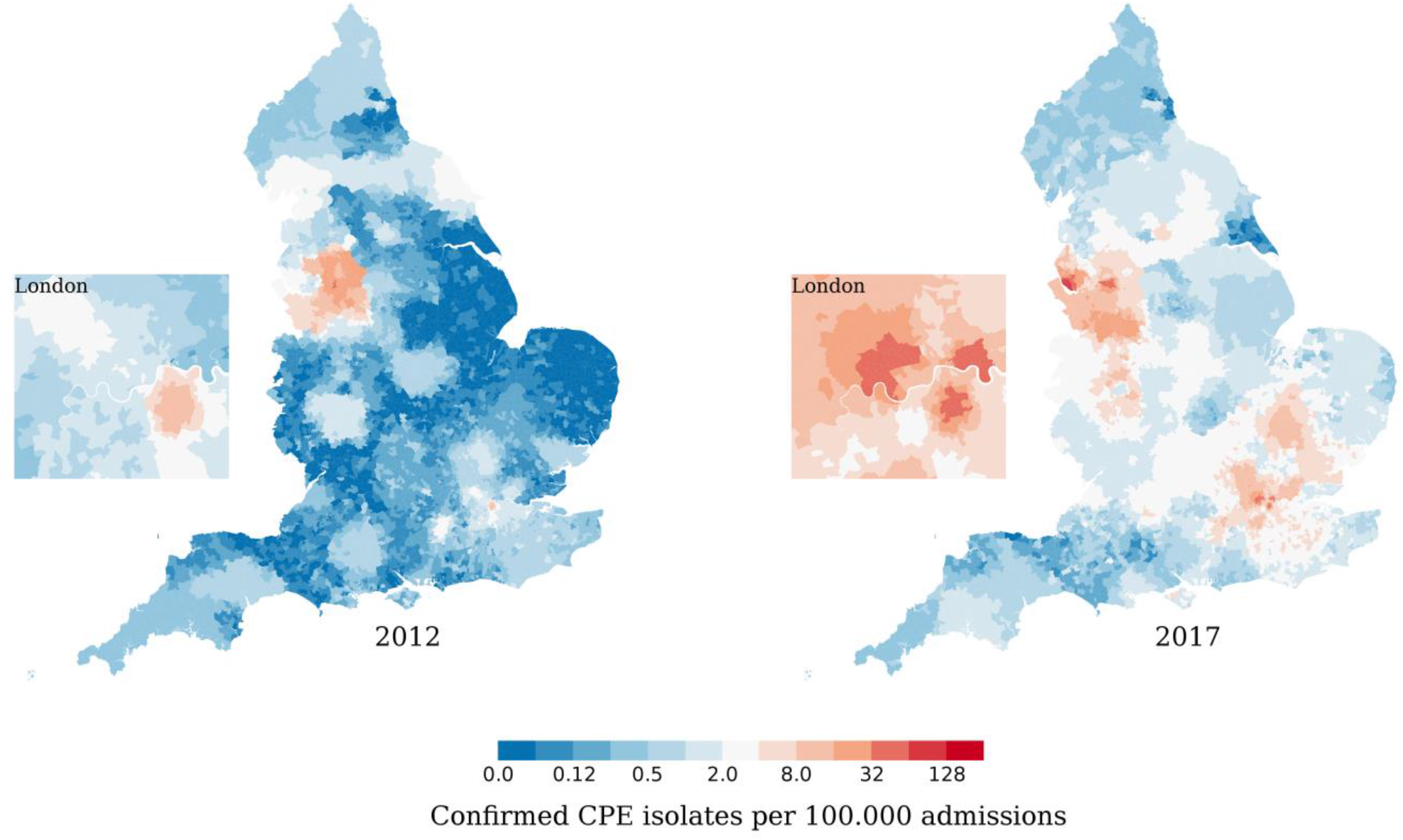
The geographical distribution of CPE in England in 2012 and 2017, based on the number of isolates confirmed by the PHE reference laboratory, as described in Donker et al [9]. The actual CPE prevalence is likely much higher, because not all locally found CPE isolates are sent to the national reference laboratory.

The increase in CPE incidence in all hospitals poses a potential problem for the control strategies described in the CPE toolkit, because the absolute number of cases from a hospital’s own local or regional population may start to outnumber the absolute number of imported cases from a ‘problem’ hospital, even if prevalence is much higher in the ‘problem’ hospital, and in the patients that have previously been admitted to it. This will reduce the effectiveness of import screening, as this strategy is purely aimed at preventing introductions. Although screening incoming patients from a problem hospital may still be the most efficient way of identifying colonised patients (from the point of view of case yield per screening effort), these cases contribute only a minority of the transmission pressure in the new hospital from this point onwards. The moment that imported cases from other hospitals become outnumbered by cases readmitted from the same/regional hospitals can be seen as a tipping point, after which hospitals need to find more effective control strategies than import screening.

To correctly assess the largest CPE risks to their hospital and their patients, hospitals thus need to be aware of the current CPE prevalence and distribution in both their own and other hospitals’ patient populations, as the correct type of control efforts against CPE crucially depends on the relative difference between these prevalences, together with the flows of patients over the hospital network. Import screening is only effective if the importation tipping point has not been reached, while control efforts aimed at reducing within-hospital transmission, such as case-finding or contact tracing, may only be effective if relatively few cases are being imported.

## When is the tipping point reached?

Hospitals can determine whether the importation tipping point has been reached using information about the number of patients received from other hospitals, the number of patients readmitted from their own hospital, the CPE prevalence among patients coming from other hospitals, and the CPE prevalence among their own readmitted patients. The CPE prevalences can be obtained by each hospital individually, by performing point-prevalence surveys among patients who had a recent previous hospital admission (including to the same hospital), while splitting the results by hospital of previous discharge. Information about the number of shared patients with other hospitals could, technically, be gathered by individual hospitals, but is often also available in centralised databases, such as the NHS Hospital-Episode Statistics (HES).

Using data presented in a previous study [9], we calculated the percentage of patients coming from other hospitals for all acute care hospital trusts in England (presented in table S1). Together with prevalence estimates from point-prevalence surveys, these can be used to calculate the percentage of expected imported cases coming from other hospitals (vs one’s own hospital) to determine if the tipping point (of when your own hospital population poses a greater risk than other ‘problem’ hospitals) has been reached. To do this, the expected number of introductions from other hospitals (calculated as the prevalence multiplied by the number of received patients) needs to be divided by the total number of expected introductions. To offer a visual aid for this process, we created a nomogram that can be used to calculate the same percentage (Figure 2 & S1).

**Figure 2).**
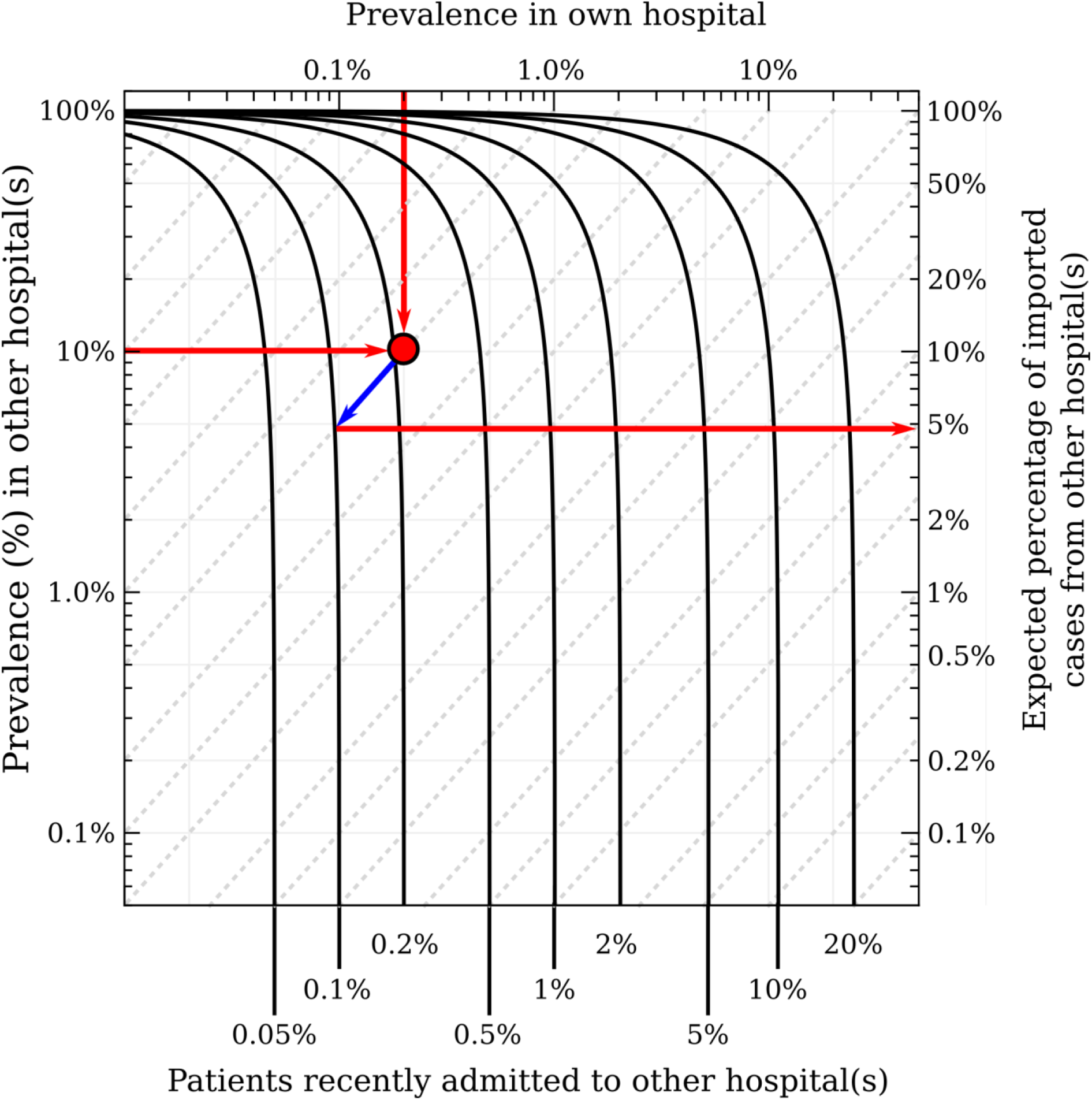
Nomogram for calculating the expected percentage of imported CPE (or any other AMR colonising pathogen) cases coming from another hospital, given the prevalence within the focal hospital, prevalence in the other hospital(s), and the percentage of readmitted patients previously admitted to the other hospital. After plotting both prevalence percentages, follow the dotted diagonal lines until the intersection with the curve for the correct percentage of readmissions that occur in patients previously discharged from the other hospital(s) (black lines); this point gives the percentage of imported cases from the other hospital on the right y-axis. Example shows a ‘self’ prevalence of 0.2%, an ‘other’ prevalence of 10%, with 0.1% of patient readmissions coming from the other hospital(s).

Using the Oxford University Hospitals Trust (OUH) as an example, we see that 89.1% of all readmitted patients have previously been admitted to the OUH itself (Table S1), while 6.5% came from other hospitals in the region, and 4.4% came from outside the region (with 0.1% of all readmitted patients coming from the Manchester region). Hypothetically, if the prevalence within the OUH’s patient population is 0.2% and the prevalence within patients discharged from any hospital in the Manchester region is 10%, just 4.8% of imported cases into the OUH would originate from the Manchester region. Even if all patients discharged from the hospitals in the Manchester region were CPE-positive (100% prevalence), they would not contribute more introductions to the OUH than patients coming from the OUH itself.

This example illustrates how the importation tipping point is reached when the prevalence within the focal hospital is relatively low. The number of shared patients from other hospitals, in particular from those hospitals further away, is diluted by the patients readmitted from the hospital itself. In general, the number of patients received from all other hospitals in the region is about 10-15% of all readmitted patients, while the total number of patients received from all hospitals outside the region is 1-3% of all readmitted patients. The tipping point is therefore often already reached when the within-hospital prevalence is two orders of magnitude lower than the prevalence in all hospitals outside the region. Similarly, the within-hospital prevalence can be 5-10-fold lower than the prevalence in all hospitals within the region to reach it.

## Discussion

Import screening to prevent introductions of CPE into hospitals is only effective at the start of an epidemic, when the number of imported cases still clearly outweigh the number of cases from the hospital itself. With the increase of CPE throughout England, it becomes more likely that import screening will no longer be a viable strategy for the majority of hospitals in the near future (if not already). In these cases, hospitals need to address the transmission of CPE within their own wards to reduce CPE prevalence, and adjust the definition of high-risk patients such that a higher proportion of the cases from its own population are identified whilst not overburdening control efforts by screening too many patients. Potential intervention strategies could in such a case still include admission screening, as long as this is no longer solely focussed at preventing introductions from other hospitals.

There is thus an immediate need to understand the transmission dynamics of CPE within hospitals to enable the design and implementation of successful control strategies. Because outbreaks of CPE are not strictly clonal, due to horizontal gene transfer, it is hard to reconstruct transmission chains through genetic relatedness, impeding our understanding of their epidemiology. Furthermore, environmental contamination may well play a role in the transmission of CPE in hospital wards [11], which might create long-term environmental reservoirs, resulting in indirect transmission events between patients who did not share time on a ward, complicating contact tracing.

Although preventing importation of CPE is not effective for CPE endemic hospitals, the effects of patient sharing between hospitals still need to be taken into account in the design of control efforts. In particular, information sharing about a patient’s CPE colonisation status, either through centralised systems or patient-held information [12], could prove essential for a timely response when these known colonised patients are admitted to another hospital. Actively tracking the admissions of CPE positive patients, through regional coordination, may reduce the need to test all patients coming from other hospitals, thus reducing unnecessary screening and isolation, and freeing up resources.

By recognising that the effectiveness of intervention strategies depends on the phase of the epidemic, we can move towards CPE control efforts that are appropriate for the current situation. Moving away from strategies based on preventing introductions from ‘problem’ hospitals when these are no longer the most effective approach can help focus much needed resources at interventions that are expected to be more effective. To enable hospitals to determine their best strategy, the CPE situation both inside and outside each hospital needs to be monitored continuously. This information will, together with knowledge about the structure of the healthcare network, help hospitals switch control strategies at the right moment.

## Supplementary Information

Table S1) The percentage of patients coming from other hospitals for all acute care hospital trusts in England

Figure S1) Nomogram for calculating the expected percentage of imported CPE (or any other AMR colonising pathogen) cases coming from another hospital. As presented in figure 2, without example data.

## Author contribution

TD, ASW and JVR drafted the manuscript, and all authors participated in writing the manuscript. All authors read and approved the final manuscript.

## Acknowledgement

None

