## Supplementary figures and images for "When do big problems far away become smaller than the problems closer to home"

### Figure S1

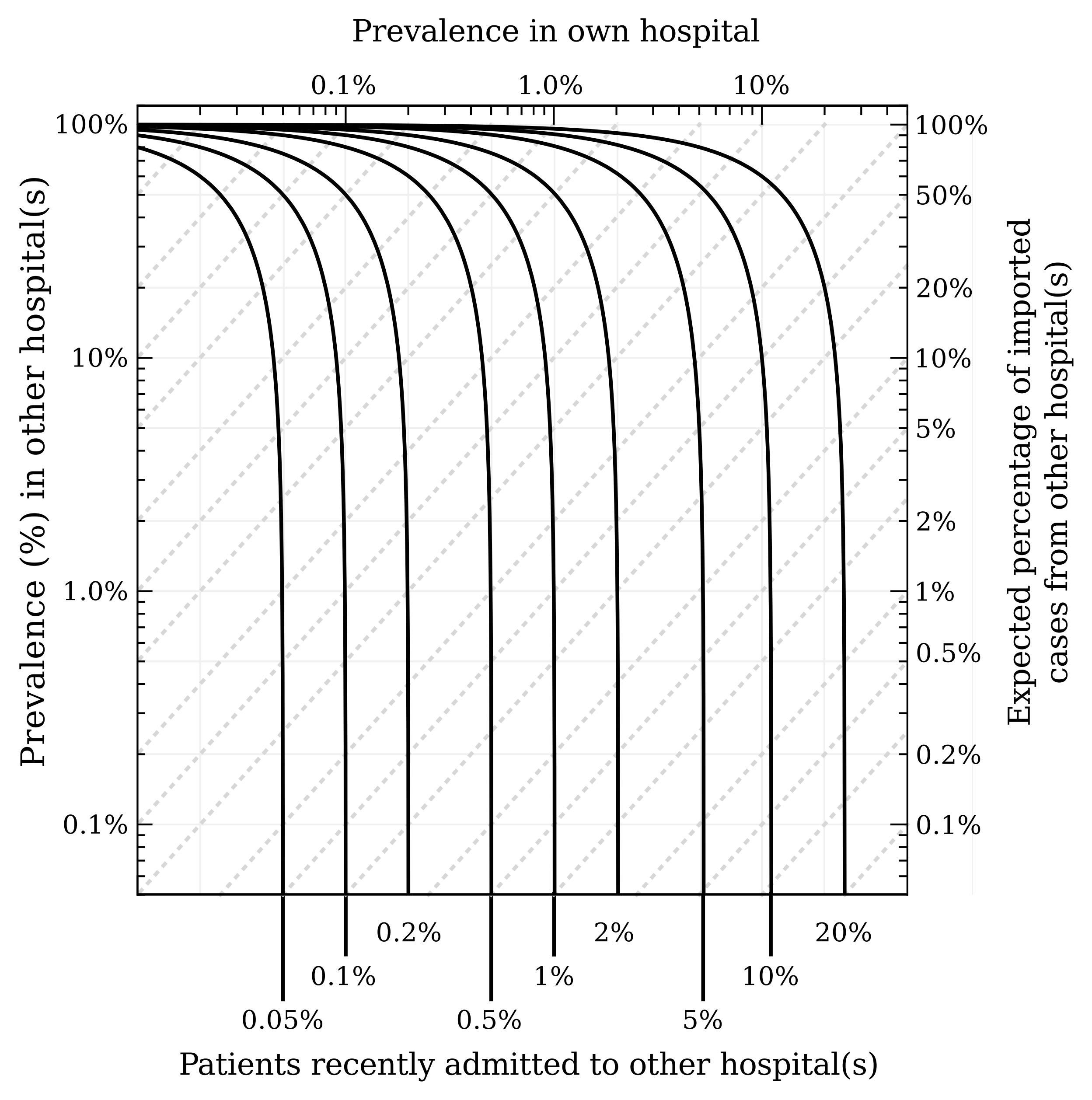
